# Evolutionary genetics of *Drosophila melanogaster* immunity: role of the X chromosome and sex-specific dominance

**DOI:** 10.1101/2020.06.03.131706

**Authors:** Manas Geeta Arun, Amisha Agarwala, Jigisha, Mayank Kashyap, Saudamini Venkatesan, Tejinder Singh Chechi, Zeeshan Ali Syed, Vanika Gupta, Nagaraj Guru Prasad

## Abstract

Intralocus Sexual Conflict (IaSC) ensues when males and females of the same species experience divergent selection on shared traits. A large number of traits have been implicated in IaSC and there is growing evidence for sexual antagonism associated with immunity. X chromosomes are thought to be hotspots of sexually antagonistic genetic variation and have been shown to harbour substantial immunity-related genetic variation.

Here, using interpopulation crosses and cytogenetic cloning, we investigated the role of the X chromosome in improved immune response of laboratory populations of the fruit-fly *Drosophila melanogaster* selected against systemic infection by *Pseudomonas entomophila*.

We could not detect any contribution of the X chromosome in the evolved immune response of our selected populations. However, we found strong evidence of sex-specific dominance related to immunity in our populations. Our results indicate that alleles that confer a superior immune response to the selected populations are, on average, partially dominant in females but partially recessive in males.

We argue that sex-specific dominance over immunity evolved as a by-product of sexually antagonistic selection in the wild ancestors of our populations. We also highlight sex-specific dominance as a potential mechanism of sex differences in immunity, with population-level sex differences primarily driven by sex differences in heterozygotes.

## Introduction

Males and females of the same species often experience distinctly different selection pressures, resulting in evolutionary conflicts (reviewed in Schenkel et al. (2018)). Sexual conflicts have been classified as interlocus sexual conflict (IeSC) or intralocus sexual conflict (IaSC). In the present study we focus solely on the latter. Typically, IaSC results when there are different fitness optima in the two sexes for a shared trait (Bonduriansky and Chenoweth 2009). At the level of a locus this translates to a scenario where different alleles are favoured in the two sexes. IaSC is thought to be resolved with the evolution of sexual dimorphism, mediated by a variety of processes such as sex-specific modification of expression, genomic imprinting and sex-specific dominance (Spencer and Priest, 2016) to name a few. In XX-XY systems, X chromosomes are thought to be hotspots of sexually antagonistic genetic variation. Using a one locus, two allele model Rice (1984) predicted that X chromosomes are expected to be more conducive than autosomes at establishing rare alleles with sexually antagonistic effects (But see Fry (2010)). This idea does have some experimental evidence in its favour. Gibson et al. (2002) cloned 20 X chromosomes from a laboratory population of *Drosophila melanogaster* and estimated that X chromosomes harbour around 45% of total fitness-related genetic variation and 97% of sexually antagonistic genetic variation. However, Ruzicka et al. (2019) could not detect a significant role of the X-chromosome in a genome-wide association study (GWAS) of sexual antagonism on a set of 202 hemi-genomes sampled from a similar base-population of *D. melanogaster.*

Immunity is an ideal trait to investigate the consequences of IaSC for at least three reasons. First, immunity is one of the traits for which differences in the two sexes have received considerable attention. In vertebrates, females consistently tend to have superior immune function relative to males (Zuk and McKean 1996; Poulin 2002). An evolutionary explanation for this trend, the immunocompetence handicap hypothesis (ICHH)(Karter and Folstad 1992), links immunity with reproduction. The ICHH argues that the immunity of males is suppressed by the action of androgens that are otherwise crucial to maintain secondary sexual characters. Evidence for ICHH is plentiful in vertebrates (reviewed in Rolff (2002)). Roved et al. (2017) argue that such hormone-mediated sex differences in immunity have the potential to lead to sexually antagonistic selection. Similar immune-endocrine interactions have been reported in invertebrates as well (Stoehr and Kokko 2006; Schwenke and Lazzaro 2017) which were previously thought to lack them. Alternative hypotheses involving trade-offs between male immunity and male ornamentation(Sheldon and Verhulst 1996) or the Bateman Principle (Rolff 2002) (but see (Stoehr and Kokko 2006)) have also been proposed. However, empirical evidence for sex differences in invertebrate immunity is equivocal. Sheridan et al. (2000), in a meta-analysis of parasite infection in arthropods, did not observe any overall sex-bias in the prevalence of infection, whereas Nunn et al. (2009) reported a female-bias in phenoloxidase activity of insects. In a recent meta-analysis, Kelly et al. (2018) reported an overall bias towards females (which was stronger in insects) but pointed out that effect sizes of this bias were small.

Second, there are numerous studies highlighting the intricate link between immunity and various aspects of reproduction. McKean and Nunney (2005) showed using *D. melanogaster* that males and females plastically modulate their investment in reproduction, depending on the availability of fitness-limiting resource, leading to sex-specific effects on immunity. Conversely, pathogenic infection has also been shown to have a sex-specific effect on reproductive fitness in *D. melanogaster* (Imroze and Prasad 2011; Nystrand and Dowling 2014). This link between immunity and reproduction has further been reinforced by numerous studies that have investigated the effect mating has on the immune response of males as well as females (McKean and Nunney 2002; Rolff and Siva-Jothy 2002; Fedorka and Zuk 2005; Kelly and Jennions 2009; Short and Lazzaro 2010; Gupta et al. 2013; Syed et al. 2020) and those investigating trade-offs between immunity and reproduction (reviewed in (Schwenke et al. 2015)).

Third, there is mounting direct evidence of IaSC related to immunity in a wide variety of taxa. Some workers have documented phenotypic evidence for sexual antagonism for immunity. Svensson et al. (2009) reported in the side-blotched lizard, *Uta stansburiana*, that an immunity phenotype that increased survival in males decreased female fitness. In a laboratory study in *D. melanogaster*, Vincent and Sharp (2014), using mutation accumulation lines, found a negative genetic correlation between the two sexes for resistance and tolerance. Using the same lines, Sharp and Vincent (2015) reported a dramatic difference in the effect of mutation accumulation on the fitness of the two sexes in presence of infection by *Pseudomonas aeruginosa* but not in the absence of the pathogen.

There is also *genetic* evidence for the sex-specific or sexually antagonistic nature of immunity. In a meta-analysis of 31 immunity-related traits in humans, Gilks et al. (2014) found that 13 traits had a higher heritability in females while 3 had a higher heritability in males, suggesting that the underlying loci were sex-dependent in their action. Genome-wide association studies (GWAS) have also identified several human SNPs that have a sex-specific influence on disease phenotypes (summarised in Gilks et al. (2014)). Hill-Burns and Clark (2009) reported in *D. melanogaster* that there is considerable immunity-related variation on the X chromosome-a hotspot for sexually antagonistic genetic variation (see below). They identified several SNPs that influenced immunity in a sexually dimorphic or antagonistic manner.

Its role in investigating immunity (Hoffmann and Reichhart 2002), sexual conflicts (see above) and life-history evolution (Prasad and Joshi 2003) makes *D. melanogaster* an ideal model organism to investigate the link between IaSc and immunity. In the present study, we used *D. melanogaster* as a model system to investigate the contribution of the X chromosome to immunity-related genetic variation and its role in adaptation to pathogenic challenge.

To that end, we used replicate laboratory populations of *D. melanogaster* selected against systemic infection by *Pseudomonas entomophila* and their respective controls. Both males and females from the selected populations have previously been shown to have evolved higher survivorship post infection relative to their counterparts from the control populations (Gupta 2016; Gupta et al. 2016)

We employed two complementary experimental approaches. First, we performed a set of interpopulation crosses, a tool previously employed to investigate the evolutionary genetics of desiccation resistance (Hoffmann and Parsons 1989), urea tolerance (Joshi et al. 1996), tolerance to chronic juvenile malnutrition (Vijendravarma and Kawecki 2013, 2015). We set up reciprocal crosses between selected populations and their respective control populations and we measured the survivorship post-infection as a proxy for the immune response of the F1 progeny.

Second, using cytogenetic cloning (Gibson et al. 2002), we sampled a set of X chromosomes from the selected and control populations and expressed them in males and females carrying the rest of the genome from the ancestral baseline population. The immune response of these flies was then assayed by measuring their survivorship post-infection.

## Methods

### I. Fly populations

All fly populations detailed below are maintained on standard banana-yeast-jaggery food unless specified otherwise.

#### A. Experimental Evolution

##### Ancestral population

Blue Ridge Baselines (BRBs) are large outbred populations of *D. melanogaster*, maintained on a 14-day discrete generation cycle, 12:12 Light:Dark regime, 25°C and 60-70% Relative Humidity. The populations have been described in detail by Singh et al. (2015).

##### Selection Regimes

We used four replicate populations (I1-4) selected against *Pseudomonas entomophila* strain L48 and their respective controls (S1-4) which have been described in detail elsewhere (Gupta 2016; Gupta et al. 2016). To summarise the selection regimes, for I populations, 2-3 day old adults are infected with *P. entomophila*, while for the S populations they are subjected to a “sham-infection” treatment (see below). In I populations, around 33% flies die over the next 96 hours. Eggs collected in the next 18 hours start the next generation. Populations with a common subscript are always handled on the same day and are related by ancestry (I1-4 and S1-4 populations were derived from the respective BRB1-4 population). They are, therefore, treated as statistical blocks. The I populations have been previously shown to have evolved a superior immune response relative to the S populations (Gupta et al. 2016).

#### B. Clone Generators (CG)

Clone generator females carry a compound X [C(1)DX *y f*] chromosome, Y chromosome, and a homozygous viable translocation of the two autosomes [T(2;3) *rdgC st in rip*^*P*^ *bw*^*D*^]. Males have an X [*sn su(b)*] chromosome, Y chromosome and the same translocated autosomes. In this system, females inherit the compound X chromosome from their mother and a Y chromosome from their father. Males inherit the Y chromosome from their mother and the X chromosome from their father. The system has been described in detail by Rice(1996). Clone generators are maintained on standard cornmeal-yeast-molasses food.

#### C. DxBRB

The DxBRB population was created by introgressing the compound X chromosome from Clone Generators into BRB_1_. This population is maintained like BRB populations.

### II. Protocol for infections/sham infections

The infection protocol involved pricking CO_2_anaesthetised flies with a needle (Minutein pin 0.1 mm, Fine Science Tools, CA) dipped in the bacterial suspension (prepared in sterile 10 mM MgSO4). For sham infections, the pricking protocol was similar, except we dipped the needle in a sterile 10 mM MgSO_4_ solution.

This study involves two distinct assays, the Hybrid Experiment and the X-Cloning Experiment, which were set up as follows:

### III. Hybrid Experiment design

Experiments were carried out between 65 and 75 generations of selection. For each experiment, we first collected eggs from I and S flies subjected to one generation of common rearing (to remove non-genetic parental effects). Ten vials of 70 eggs each were set up per population. Adult males and females that emerged from these vials were collected as virgins and were combined in Plexiglas cages to set up the following crosses of 100 pairs each.

1. I ♀ × I ♂ (II)
2. S ♀ × S ♂ (SS)
3. I ♀ × S ♂ (IS)
4. S ♀ × I ♂ (SI)

In order to generate the F1 progeny to be used in our experiments, we collected 10 vials each containing 70 eggs for every cross. On the 12^th^-day post egg collection, for every cross, we set up three replicate cages, each containing 50 males and 50 females infected with *P. entomophila* (OD_600_=1.5). Additionally, for each cross, we also set up a cage containing 50 male and 50 females that were sham-infected. We monitored mortality over the next four days. Food plates in the cages were replaced with fresh ones two days after the cages were set up.

### IV. X-Cloning Experiment

#### Cloning of the X chromosome

After 160 generations of selection, 30 X chromosomes were randomly sampled from each I and S population, each used to create a single X chromosome line.

In order to express the X chromosome from I and S populations in a neutral chromosomal background, the following crosses were set up over 4 generations (See Figure S3 for a detailed schematic):

1. I/S♂ × CG♀ → Progeny1_(Males)_ + other progeny
2. Progeny1♂ × DxBRB♀ → Progeny2_(Brown-eyed Males)_+ other progeny
3. Progeny2♂ × DxBRB♀ → Progeny3_(Red-eyed Males)_+ other progeny
4. Progeny3♂ × DxBRB♀ → Experimental flies_(Males)_+ other progeny Progeny3♂ × BRB_1_♀ → Experimental flies_(Females)_+ other progeny

Males from I and S populations were crossed to Clone Generator females. Males resulting from this cross were crossed to virgin DxBRB females. The brown-eyed males resulting from that cross were again crossed to virgin DxBRB females. In order to generate male and female flies to be used in the survivorship assay, red-eyed males resulting from the previous cross were crossed to virgin DxBRB and BRB females respectively. The experimental flies carry the X chromosome from I or S populations, but the rest of the genome is from BRB (other than the unmanipulated fourth chromosome.) A single vial per X-line was maintained for all crosses. Egg densities were maintained such that there were 70 viable eggs per vial. Crosses were set on the 12^th^-day post egg collection.

#### Survivorship Assay

For the survivorship assay, 20 X-lines per population were randomly selected. On the 11^th^-day post egg-collection (of the 4^th^ cross), flies were sorted into same sex vials (10 per vial). On the 12^th^ day, flies were infected with *P. entomophila* (OD_600_=1) and transferred to fresh vials. 3 vials per X-line (8 flies per vial) were set up for the infected treatment. We also set up one vial per X-line of sham controls. Mortality was monitored over a 96-hour period. The surviving flies were transferred to fresh food vials after 2 days.

We use survivorship post-infection as a read-out of immunity as I populations have previously been shown to have evolved higher survivorships post infection relative to S populations during the course of adapting to systemic infection by *P. entomophila* (Gupta et al. 2016). *P. entomophila*, originally isolated from a wild *D. melanogaster* female, is a widely used model pathogen for immunity studies on *D. melanogaster* (Dieppois et al. 2015). This makes survivorship post infection by *P. entomophila* an ecologically relevant read-out of immunity. Furthermore, it has been used as a proxy for immunity in other studies employing experimental evolution of *D. melanogaster* immunity (Martins et al. 2013; Faria et al. 2015). In *D. melanogaster*, survivorship post-infection is also fairly strongly correlated with other immune read-outs such as the ability to restrict bacterial growth (measured using colony forming units or CFUs) and expression of anti-microbial peptides (AMPs) (Schwenke and Lazzaro 2017).

#### Statistical Analysis

In neither of our experiments was there any mortality in the sham control treatment. Therefore, in subsequent analyses data for the sham treatment was excluded. All analyses were performed on R (version 3.5.3).

##### A) Hybrid Experiment

We performed three different analyses for this experiment.

First, for each combination of cross, sex, infector and block we calculated the proportion of flies alive at the end of the 96-hour observation window. Using this proportion as the unit of analysis we fit the following linear mixed effects model using the R packages “lme4” and “lmerTest”:

Proportion Survivorship ∼ Cross + Sex + Cross:Sex + (1 | Block) + (1 | Sex:Block) + (1 | Cross:Block) + (1|Infector).

Post-hoc comparisons using Tukey’s HSD were implemented with the R package “emmeans”.

Second, we used the R package “coxme” and fit the following Cox’s proportional hazards model:

Time to death ∼ Cross + Sex + Cross:Sex + (1 | Block/Sex/Cross) + (1 | Infector)

Third, we used the R package lme4 to fit the following model using logistic regression on the status (dead or alive) of each individual fly at the end of the 96-hour observation window.

Status ∼ Cross + Sex + Cross:Sex + (1 | Infector) + (1 | Block)

Our results (Table S1, Figure 1) suggested that differences in the two sexes were primarily driven by differences between the two sexes within the hybrid crosses (IS and SI) but not within the parental crosses (II and SS). Therefore, in order to investigate the possibility of sex-specific dominance, we also calculated an estimate for the dominance coefficient for proportion survivorship for both sexes. Since neither of our analyses could distinguish between IS and SI (see results), we used the average proportion survivorship of IS and SI of the ‘n’th Block as the average “heterozygote” proportion survivorship for that block. We used the following expression to calculate the dominance coefficient:

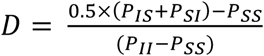

where “*P*_*AB*_*”* stands for proportion survivorship of the cross “AB”. (Ewens 2004, adapted from fitness scheme 1.25b, Section 1.4)

**Figure 1.**
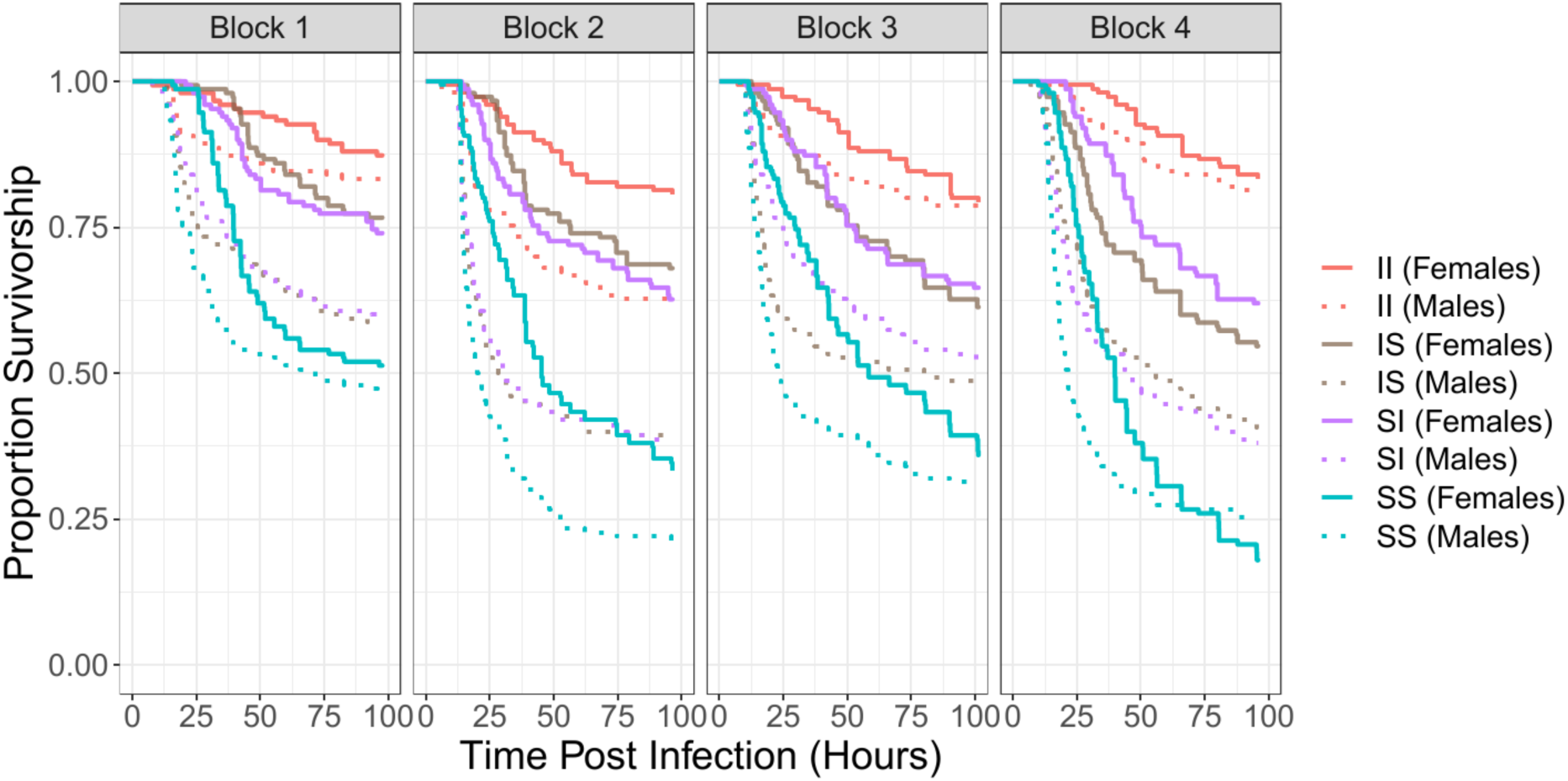
Effect of the cross and sex on survivorship post-infection in the Hybrid Experiment. The curves show survival of the F_1_ progeny as a function of time. The first letter indicates the maternal selection regime and the second, the paternal.

To test for additivity, we performed separate t tests for males and females.

##### B) X-cloning Experiment

We performed four different analyses for this data.

We calculated proportion survivorship at the end of the 96-hour observation window for each combination of block, selection regime, sex and X chromosome line. For this purpose, we pooled the data from the three vials for each X chromosome line. We then fit the following linear mixed-effects model using the R package lme4:

Y ∼ SelectionRegime + Sex + SelectionRegime:Sex + (1 | Block) + (1 | X-line:SelectionRegime:Block)

We calculated the median time to death for each vial and fit the following linear mixed-effects model using lme4:

Y ∼ SelectionRegime + Sex + SelectionRegime:Sex + (1 | Infector) + (1 | Block) + (1 | X-line:SelectionRegime:Block)

We fit the following logistic regression on the status (dead or alive) of each fly at the end of the 96-hour observation window:

Status ∼ Selection Regime + Sex + SelectionRegime:Sex + (1 | Infector) + (1 | Block) + (1 | X-line:Block:SelectionRegime)

We fit the following cox’s proportional hazards model:

Time to Death ∼ SelectionRegime+ Sex + SelectionRegime:Sex +(1 | Block/SelectionRegime/Xline) + (1 | Infector)

We also fit the following cox’s proportional hazards model separately for each block:

Time to Death ∼ SelectionRegime + Sex + SelectionRegime:Sex + (1 | Infector) + (1 | X-line/SelectionRegime)

Additionally, we calculated the average median time to death and proportion survivorship for each X-line in both the sexes. For these two read-outs of immunity, we calculated the correlation between male and female immunity, separately for each combination of selection regime and block.

## Results

### A) Hybrid Experiment

In our linear mixed-effects model, we found a significant effect of Sex, Cross as well as their interaction (Table 1A). The effect of Sex was primarily, if not entirely, a result of the differences between the two sexes within the hybrid crosses. A post-hoc Tukey’s HSD suggested that in the case of II and SS the two sexes were not significantly different, but in the two hybrid crosses, males fared significantly worse than females (Table S1). Furthermore, in both sexes, the II cross had the highest survivorship, followed by IS and SI, which were not significantly different from each other. SS had the worst survivorship (Figure 1, Figure S1).

**Table 1.**
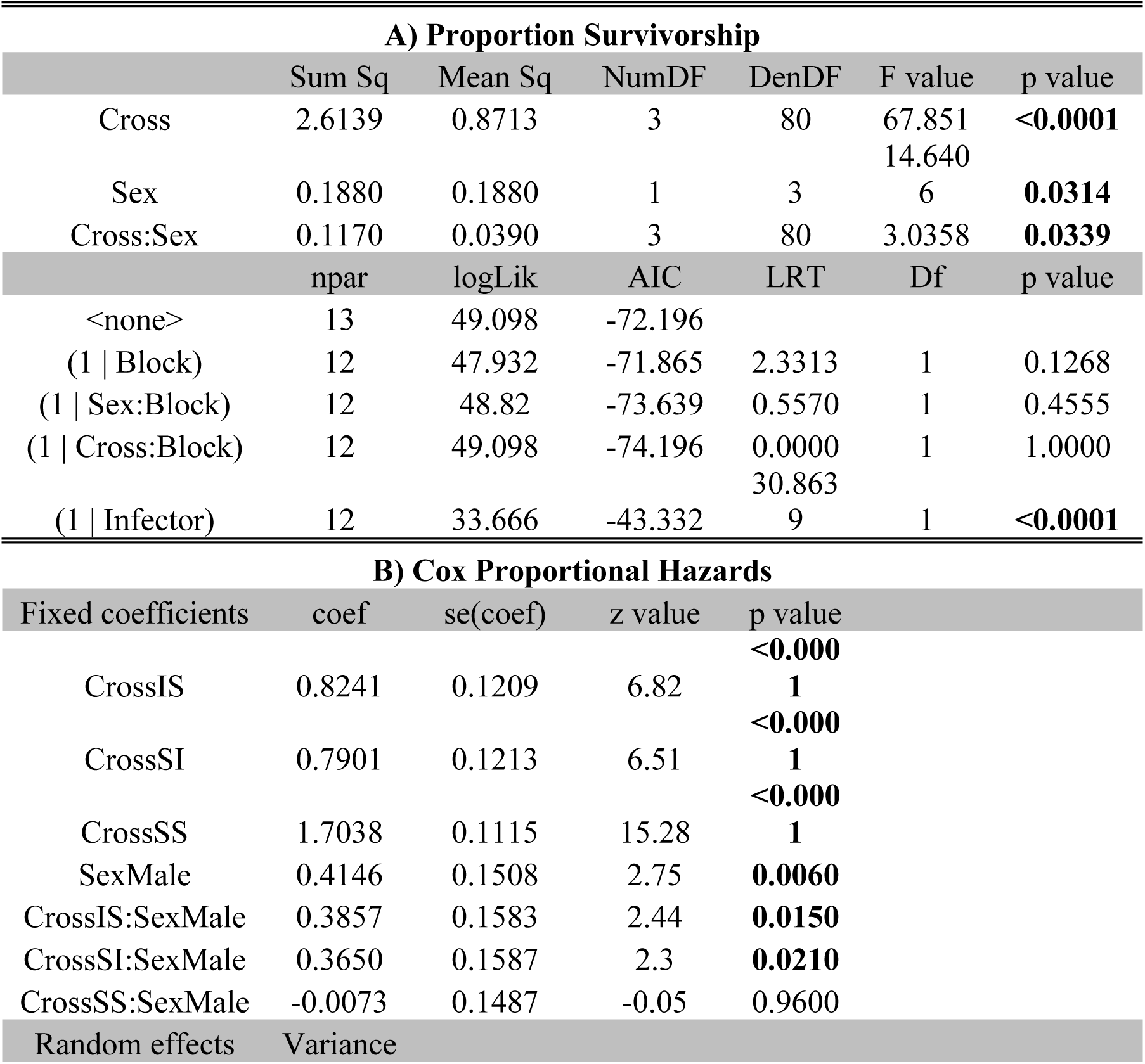
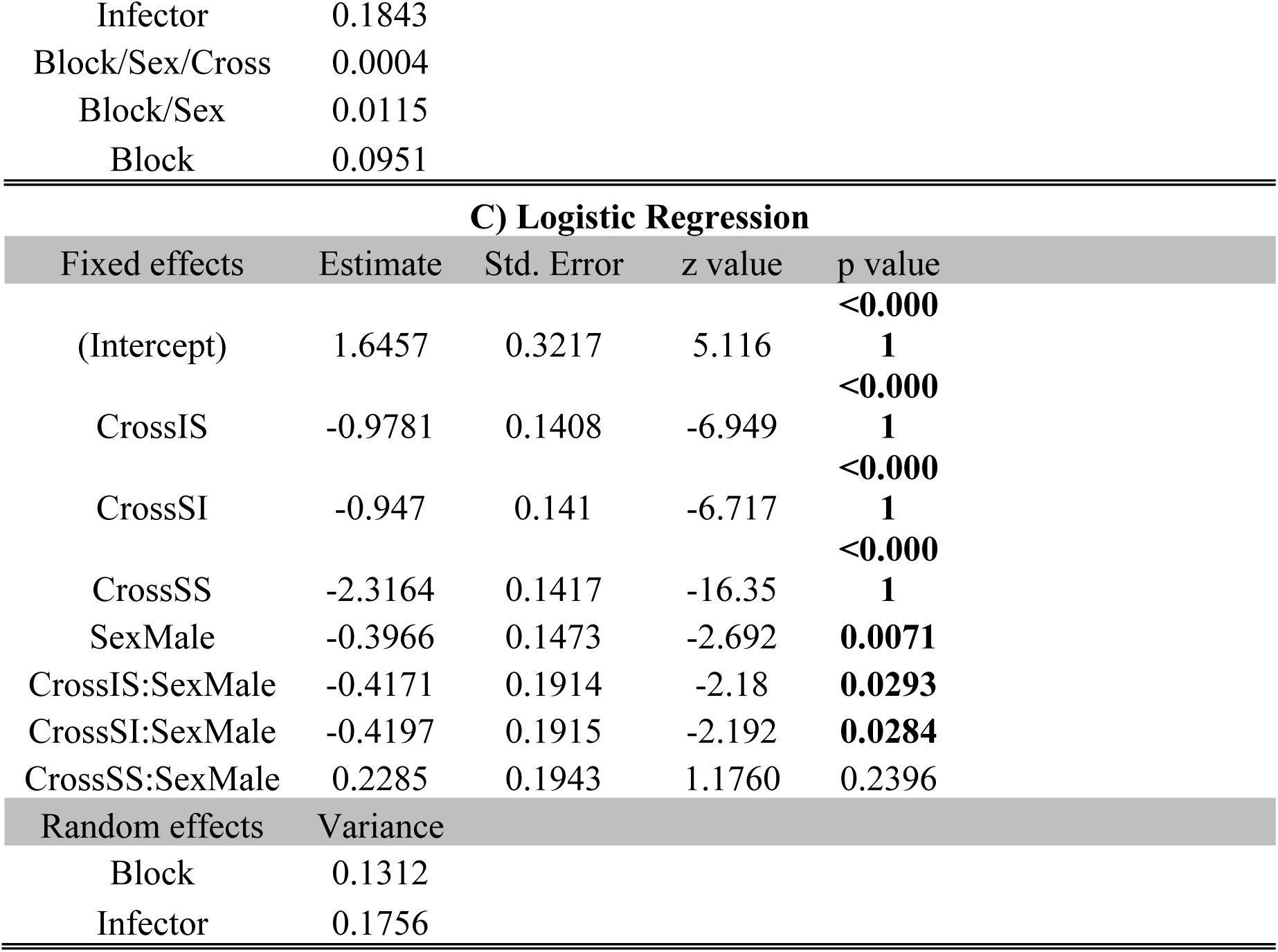
Summary of Hybrid Experiment results. A) Linear mixed effects model of proportion survivorship, B) Cox Proportional Hazards model of survivorship post-infection, C) Logistic Regression

The results of our Cox’s proportional hazards model and logistic regression were qualitatively similar (Table 1B, C). Both analyses suggested that the two hybrid crosses were similar to each other but worse than the II cross. The SS cross had the worst survivorship. Overall, females had higher survivorship compared to males. In both analyses the coefficients associated with CrossIS:SexMale and CrossSI:SexMale were significantly different from 0, suggesting that the pattern of sex differences in these crosses was different from the pattern of sex differences in the II cross. At the same time the coefficient associated with CrossSS:SexMale was not significantly different from 0 suggesting that the pattern of sex differences in the SS cross was similar to the II cross. This clearly suggests that the significant effect of Sex was mainly a result of sex differences within the two hybrid crosses, SI and IS (Figure 1).

The dominance coefficient for females (mean = 0.6459, standard deviation = 0.0298) was significantly greater than 0.5 (p = 0.0019), while the same for males (mean = 0.3526, standard deviation = 0.0747) was significantly less than 0.5 (p = 0.0271).

### B) X-Cloning Experiment

In our logistic regression and the linear mixed effects model for proportion survivorship, we found a significant effect of sex, with males having marginally higher survivorships than females, while there was no effect of selection regime or its interaction with sex (Table 2A,C, Figure S2A). The linear mixed effects model for median time to death failed to detect any effect of selection regime, sex or their interaction (Table 2B, Figure S2B).

**Table 2.** Summary of X-Cloning Experiment results. A) Logistic Regression, Linear mixed effects model of B) median time to death and C) proportion survivorship, D) Cox Proportional Hazards model of survivorship post-infection

| A) Logistic Regression |  |  |  |  |
| --- | --- | --- | --- | --- |
| Fixed Effects | Estimate | Std. Error | z value | p value |
| (Intercept) | -3.2541 | 0.2689 | -12.1 | **<0.0001** |
| SelectionS | -0.2443 | 0.1857 | -1.315 | 0.1884 |
| Sexmale | 0.4386 | 0.1401 | 3.13 | **0.0018** |
| SelectionS:Sexmale | 0.1502 | 0.2069 | 0.726 | 0.4678 |
| Random Effects | Variance |  |  |  |
| Xline:Block:Selection | 0.3050 |  |  |  |
| Block | 0.0566 |  |  |  |
| Infector | 0.1235 |  |  |  |

| B) Median Time to Death |  |  |  |  |  |  |
| --- | --- | --- | --- | --- | --- | --- |
|  | Sum Sq | Mean Sq | NumDF | DenDF | F value | p value |

| Selection | 439354 | 439354 | 1 | 156.13 | 1.6122 | 0.2061 |
| --- | --- | --- | --- | --- | --- | --- |
| Sex | 11070 | 11070 | 1 | 801.11 | 0.0406 | 0.8403 |
| Selection:Sex | 387 | 387 | 1 | 801.11 | 0.0014 | 0.9699 |
|  | npar | logLik | AIC | LRT | Df | p value |
| <none> | 8 | -7360.8 | 14738 |  |  |  |
| (1 \| Infector) | 7 | -7374.8 | 14764 | 28.0106 | 1 | **<0.0001** |
| (1 \| Block) | 7 | -7373.9 | 14762 | 26.2017 | 1 | **<0.0001** |
| (1 \| Xline:Selection:Block) | 7 | -7360.9 | 14736 | 0.1795 | 1 | 0.6718 |

C) Proportion Survivorship
|  | Sum Sq | Mean Sq | NumDF | DenDF | F value | p value |
| --- | --- | --- | --- | --- | --- | --- |
| Selection | 0.0054 | 0.005367 | 1 | 155.07 | 1.4284 | 0.2338 |
| Sex | 0.0476 | 0.04758 | 1 | 158.01 | 12.6633 | **0.0005** |
| Selection:Sex | 0.0002 | 0.000159 | 1 | 158.01 | 0.0423 | 0.8374 |
|  | npar | logLik | AIC | LRT | Df | p value |
| <none> | 7 | 422.33 | -830.66 |  |  |  |
| (1 \| Xline:Block:Selection) | 6 | 422.33 | -832.66 | 0.0029 | 1 | 0.9572 |
| (1 \| Block) | 6 | 419.92 | -827.84 | 4.8251 | 1 | **0.0281** |

D) Cox Proportional Hazards
| Fixed Coefficients | coef | se(coef) | z value | p value |
| --- | --- | --- | --- | --- |
| SelectionS | 0.0583 | 0.0558 | 1.05 | 0.3000 |
| Sexmale | 0.0131 | 0.0342 | 0.38 | 0.7000 |
| SelectionS:Sexmale | 0.0293 | 0.0483 | 0.61 | 0.5400 |
| Random effects | Variance |  |  |  |
| Block/Selection/Xline | 0.0777 |  |  |  |
| Block/Selection | <0.0001 |  |  |  |
| Block | 0.0827 |  |  |  |
| Infector | 0.0368 |  |  |  |

Neither of our linear mixed effects models (proportion survivorship or median time to death) could detect an effect of the X chromosome line.

In our Cox’s proportional hazards model that incorporates block as a random factor (Table 2D, Figure 2), we did not detect a significant effect of selection regime, sex or their interaction.

**Figure 2.**
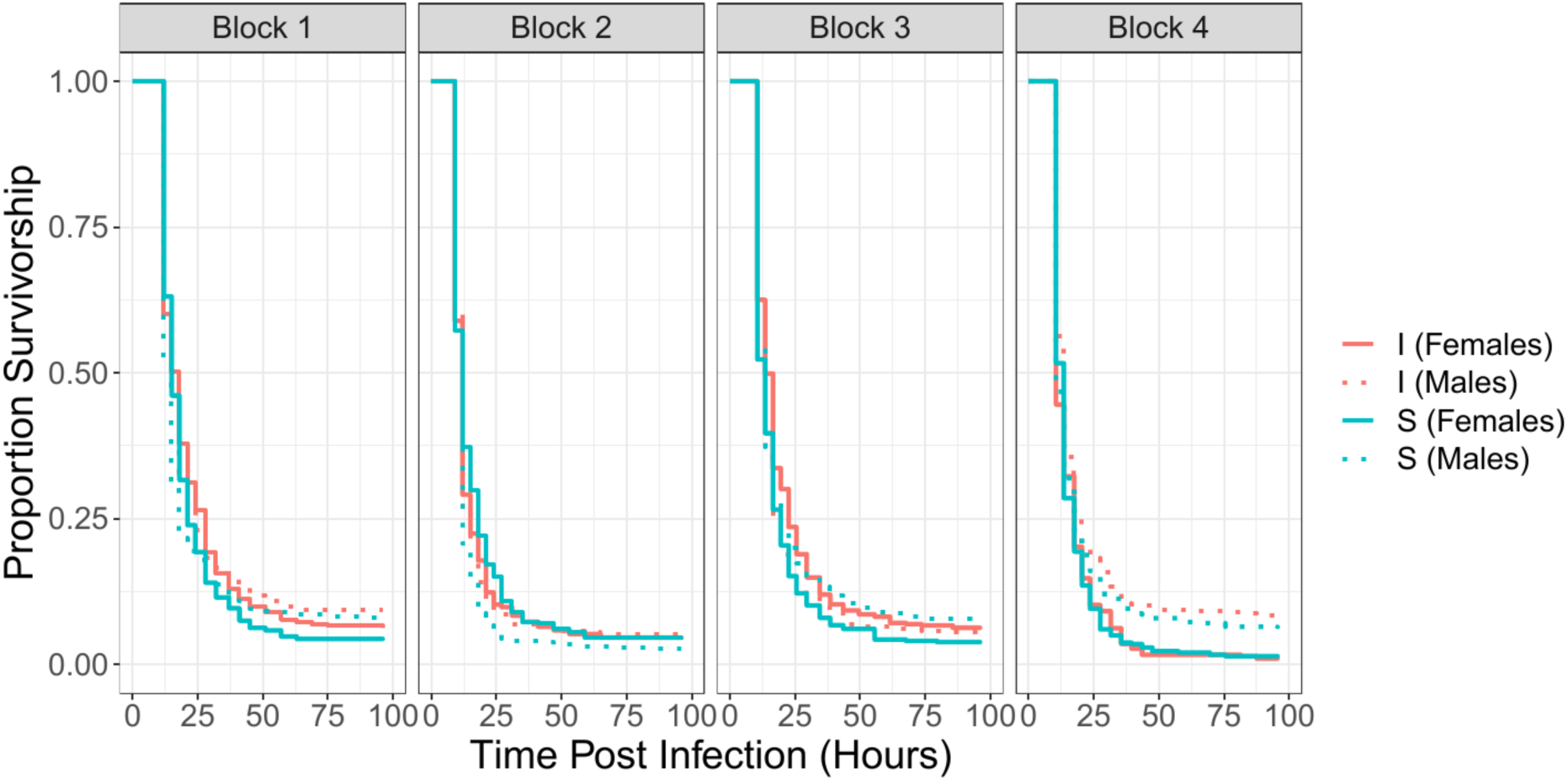
Effect of Selection Regime and sex on survivorship post-infection in the X-Cloning Experiment. The curves show survival as a function of time for I females (solid orange), I males (dashed orange), S females (solid blue) and S males (dashed blue). I and S flies carry the X chromosome from the respective selection regime but share the rest of the genome, which comes from a neutral baseline population.

For our separate blockwise Cox’s proportional hazards models we could not detect an effect of selection regime, sex or their interaction in three of the four blocks (Table S3, Figure 2). In Block 3, we found a significant effect of selection regime, sex and their interaction. Females carrying X chromosomes from the I3 population had a slightly higher survivorship than females carrying X chromosomes from the S3 population, while males carrying S3 X chromosomes had marginally higher survivorship than males carrying I3 X chromosomes. However, as is apparent in Figure 1, the magnitude of these differences was fairly small.

In seven of the eight selection regime × block combinations we did not detect a significant correlation between male and female proportion survivorship or median time to death using Spearman’s rank correlation or the linear model (Table S2A,B). In I2 we find a significant but weak (R^2^=0.353) correlation between male and female proportion survivorship in the linear model (Table S2B).

## Discussion

We set up crosses between replicate populations of *Drosophila melanogaster* selected against infection by *Pseudomonas entomophila* with their respective controls. Subsequently, we measured the immune response of the F1 progeny. We also isolated X chromosomes from the selected and control populations and measured their contribution to the immune response of flies carrying common ancestral autosomes. Here, we discuss two key findings of our experiment.

### 1. Sex-specific dominance

A rather unexpected finding of our study was that the alleles that conferred improved immunity to I males and females had different dominance coefficients in the two sexes. The survivorship of males was much worse than females in the two hybrid crosses (IS and SI) but not so in the two parental crosses (Figure 1). Furthermore, our analysis of dominance coefficients indicated that the “better-immunity” alleles on average were partially dominant in females (dominance coefficient = 0.6471), but partially recessive in males (dominance coefficient= 0.3541). To the best of our knowledge, this is the first report of sex-specific dominance in the case of any immunity-related trait.

We believe that sex-specific dominance in immunity in our populations is a signature of sexual antagonism historically experienced by the wild ancestors of our populations.

The link between sexual antagonism and sex-specific dominance has only recently begun to be investigated. Using a two-locus model, Spencer and Priest (2016) showed that a modifier allele that modulates the dominance coefficient of a sexually antagonistic locus in a sex-specific manner can indeed invade a population, leading to the evolution of sex-specific dominance coefficients. Empirical evidence for this idea is sparse. But Barson et al. (2015) were able to identify a locus in salmon which exhibits sex-specific dominance for age at maturity, a trait under sexually antagonistic selection. Grieshop and Arnqvist (2018) used a diallel cross design on isogenic lines from a population of *Callosobruchus maculates* and found strong evidence for sex-specific dominance for sexually antagonistic polymorphisms.

We speculate that sex-specific dominance for immunity-related loci in our *Drosophila* populations could be a result of a similar process. It is very unlikely that modifiers bringing about sex-specific dominance would spread over the course of the duration of our laboratory selection experiment (<100 generations). We speculate that in the wild ancestors of our laboratory populations, the alleles that conferred a superior immune response were favoured in females, while alleles that conferred a poorer immune response were favoured in males through their pleiotropic action on male fitness in other contexts (e.g. reproduction). Given the Bateman Principle, this is not an unrealistic assumption and has been invoked quite often (Rolff 2002; McKean and Nunney 2005). In fact, studies have shown that I males tend to have a poorer mating success when directly competing with S males (Venkatesan 2015). We believe that this antagonistic selection resulted in the evolution of sex-specific dominance for immunity-related alleles such that female-beneficial alleles (also the “better immunity” alleles) evolved to become more dominant in females, but less dominant in males. During the course of our study, these alleles that conferred superior immunity in both sexes, but were more dominant in females than in males, increased in frequency in the I population as a result of strong selection on immunocompetence.

Our results are important in the context of sex differences in immunity. As a result of sex-specific dominance, sex differences in immunity at a population level could arise solely through the difference in immunocompetence of the heterozygote genotype expressed in the two sexes. This, of course, would require the maintenance of heterozygotes at sufficiently high frequencies, through processes such as trade-offs between male immunocompetence and reproductive output.

### 2. No evidence of an effect of X chromosome

X chromosomes have also been predicted to be hotspots of sexually antagonistic fitness variation (Rice 1984). *D. melanogaster* X chromosomes harbour 45% of the total fitness variation and 97% of the total sexually antagonistic fitness variation (Gibson et al. 2002). Given the considerable evidence for immunity-related sexual antagonism in *D. melanogaster*, we expected the improvement in the immune response of the I populations to be a result of, largely, evolution of X-linked immunity-related loci. Differences in the X chromosomes from I and S populations could also arise as a consequence of the “Faster X Effect” (Hartl 1972; Charlesworth et al. 1987) which posits that adaptive evolution should, typically, occur faster on the X chromosomes relative to autosomes.

But contrary to our expectation, male progeny from the two hybrid crosses (IS and SI) had indistinguishable survivorships post infection suggesting that the X chromosomes from I and S populations were similar in their immune performance, at least in males. One drawback of this design, however, is that it does not take into account the Y chromosome. Apart from possessing X chromosomes from different selection regimes, IS and SI males also inherit Y chromosomes belonging to different selection regimes. While Y chromosomes are generally thought to be depauperate in genes, Kutch and Fedorka (2017) reported the presence of Y-linked variation that regulates autosomal immune function genes in *D. melanogaster*. Therefore, the Y chromosome could potentially confound our conclusion. Our findings from the X-cloning experiment, however, rule out this possibility. In neither of our analyses could we distinguish between X chromosomes from the I populations from X chromosomes from the S populations with respect to their immune response. This clearly suggests that the improvement in the immunity of the I populations did not involve loci located on the X chromosome. This apparent dearth of X-linked immunity-related loci among the loci that have evolved in the I populations is significant given that in *D. melanogaster* the X chromosome contains 20% of the total genome (Turelli and Begun 1997).

In his model Rice (1984) assumed that the dominance coefficients were identical in the two sexes. Fry (2010) showed that if one relaxes this assumption such that the male (female) beneficial allele is at least partially dominant in males (females), autosomes are better than the X chromosome in facilitating a sexually antagonistic polymorphism. In the light of our results indicating sex-specific dominance, a lack of effect of the X chromosome in our experiments is quite unsurprising. Our results are consistent with a series of recent studies using laboratory populations of *D. melanogaster* that did not find unequivocal evidence in support of the idea that the X-chromosome is a hotspot for sexually antagonistic genetic variation. Ruzicka et al. (2019) used GWAS using hemiclonal analysis and could not detect significant X-linked sexual antagonism. Abbott et al. (2020) restricted the evolution of the X-chromosome to males and found an increase in male fitness but not a corresponding decrease in female fitness expected under sexual antagonism. Lund-Hansen et al. (2020) carried out the reciprocal experiment by restricting X-chromosome evolution to females resulting in the “feminization” of body-weight and development time but not female reproductive fitness and locomotory activity, a trait previously shown to be under IaSC (Long and Rice 2007)

Our results are in stark contrast to the findings of Hill-Burns and Clark (2009), who had reported considerable immunity-related variation on the X chromosome. Our results indicate that selection for improved immune response did not result in any evolution of the X chromosome. At a more fundamental level, in our X-cloning experiment, we could not detect an effect of X-line, suggesting that there is negligible amount of X-linked immunity-related genetic variation in our populations. It must be noted, however, that Hill-Burns and Clark had used bacterial clearance ability as a measure of immunity, which may not necessarily translate to improved survival in the face of pathogenic infection.

Vincent and Sharp (2014) had found a negative genetic correlation between male and female immune components. For X chromosomes derived from 3 selected and 4 control populations, we failed to detect any such male-female correlations.

## Conclusion

Ours is among the first studies to use experimental evolution to investigate the genetic architecture of the *D. melanogaster* immune response. Our study throws light on two important aspects of *D. melanogaster* immunity genetics. Firstly, very few immunity-related loci that aid a population to adapt in the face of systemic pathogenic infection are located on the X chromosome. Secondly, ours is the first study to report evidence of sex-specific dominance in the immune response of *D. melanogaster.* Furthermore, we identify sex-specific dominance coefficients as a potential mechanism of explaining sex differences in immunity.

## Acknowledgements

VG, ZAS, MGA thank CSIR, Govt. of India for Junior and Senior Research Fellowships. TSC thanks IISER Mohali for Senior Research Fellowship. AA thanks DST, Govt. of India for the KVPY scholarship. SV MGA J MK thank DST, Govt. of India for INSPIRE-SHE. This work was funded by IISER Mohali and DBT, Govt. of India (Grant No. DBT-17-0128).

We would like to thank Sharmi Sen, Aparajita Singh and Naginder for their help during the experiments, Dr. Pierre Cornelis of the Free University of Brussels, Belgium, who provided us with the bacterial strain used here and Prof. Adam Chippindale for generously providing us with the Clone Generator flies.

