## Supplementary Information for "Evolutionary genetics of *Drosophila melanogaster* immunity: role of the X chromosome and sex-specific dominance"

**Hybrid Experiment**

**Table S1** Tukey's HSD for proportion survivorship in the Hybrid Experiment

| contrast | | estimate | SE | df | t.ratio | p value |
|---|---|---|---|---|---|---|
| II,Female | IS,Female | 0.1750 | 0.0463 | 31.9 | 3.783 | 0.0132 |
| II,Female | SI,Female | 0.1683 | 0.0463 | 31.9 | 3.639 | 0.019 |
| II,Female | SS,Female | 0.4800 | 0.0463 | 31.9 | 10.376 | <.0001 |
| II,Female | II,Male | 0.0617 | 0.0505 | 19.5 | 1.222 | 0.9157 |
| II,Female | IS,Male | 0.3617 | 0.0505 | 15.2 | 7.165 | 0.0001 |
| II,Female | SI,Male | 0.3550 | 0.0505 | 15.2 | 7.033 | 0.0001 |
| II,Female | SS,Male | 0.5150 | 0.0505 | 15.2 | 10.202 | <.0001 |
| IS,Female | SI,Female | -0.0067 | 0.0463 | 31.9 | -0.144 | 1 |
| IS,Female | SS,Female | 0.3050 | 0.0463 | 31.9 | 6.593 | <.0001 |
| IS,Female | II,Male | -0.1133 | 0.0505 | 15.2 | -2.245 | 0.3795 |
| IS,Female | IS,Male | 0.1867 | 0.0505 | 19.5 | 3.698 | 0.026 |
| IS,Female | SI,Male | 0.1800 | 0.0505 | 15.2 | 3.566 | 0.0433 |
| IS,Female | SS,Male | 0.3400 | 0.0505 | 15.2 | 6.735 | 0.0001 |
| SI,Female | SS,Female | 0.3117 | 0.0463 | 31.9 | 6.737 | <.0001 |
| SI,Female | II,Male | -0.1067 | 0.0505 | 15.2 | -2.113 | 0.4486 |
| SI,Female | IS,Male | 0.1933 | 0.0505 | 15.2 | 3.83 | 0.0265 |
| SI,Female | SI,Male | 0.1867 | 0.0505 | 19.5 | 3.698 | 0.026 |
| SI,Female | SS,Male | 0.3467 | 0.0505 | 15.2 | 6.867 | 0.0001 |
| SS,Female | II,Male | -0.4183 | 0.0505 | 15.2 | -8.287 | <.0001 |
| SS,Female | IS,Male | -0.1183 | 0.0505 | 15.2 | -2.344 | 0.3319 |
| SS,Female | SI,Male | -0.1250 | 0.0505 | 15.2 | -2.476 | 0.2748 |
| SS,Female | SS,Male | 0.0350 | 0.0505 | 19.5 | 0.693 | 0.9962 |
| II,Male | IS,Male | 0.3000 | 0.0463 | 31.9 | 6.485 | <.0001 |
| II,Male | SI,Male | 0.2933 | 0.0463 | 31.9 | 6.341 | <.0001 |
| II,Male | SS,Male | 0.4533 | 0.0463 | 31.9 | 9.799 | <.0001 |
| IS,Male | SI,Male | -0.0067 | 0.0463 | 31.9 | -0.144 | 1 |
| IS,Male | SS,Male | 0.1533 | 0.0463 | 31.9 | 3.314 | 0.042 |
| SI,Male | SS,Male | 0.1600 | 0.0463 | 31.9 | 3.459 | 0.0297 |

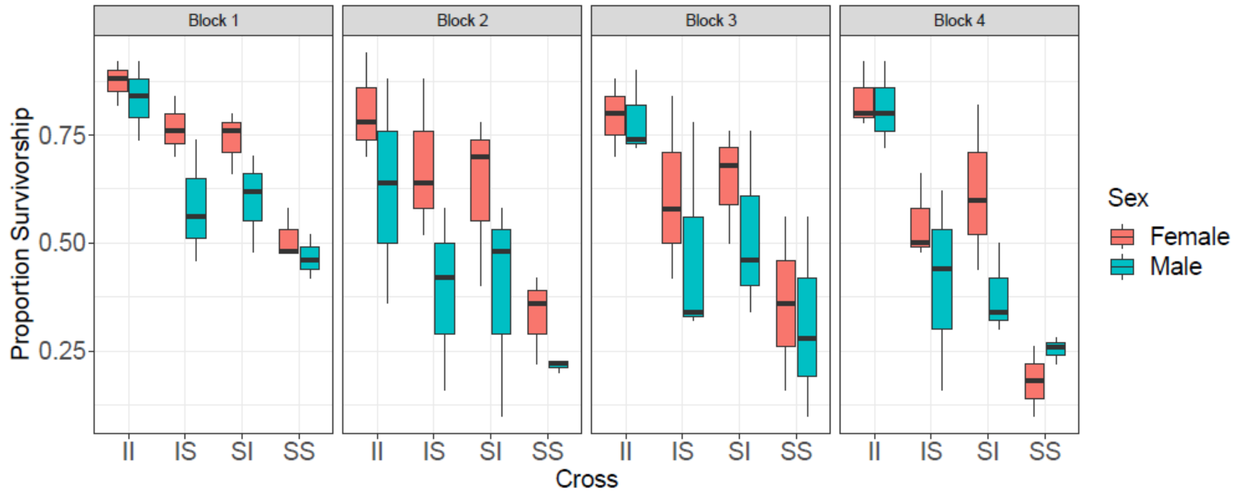


**Figure S1** Effect of cross and sex on proportion survivorship at the end of the observation window in the Hybrid Experiment.

### X Cloning Experiment

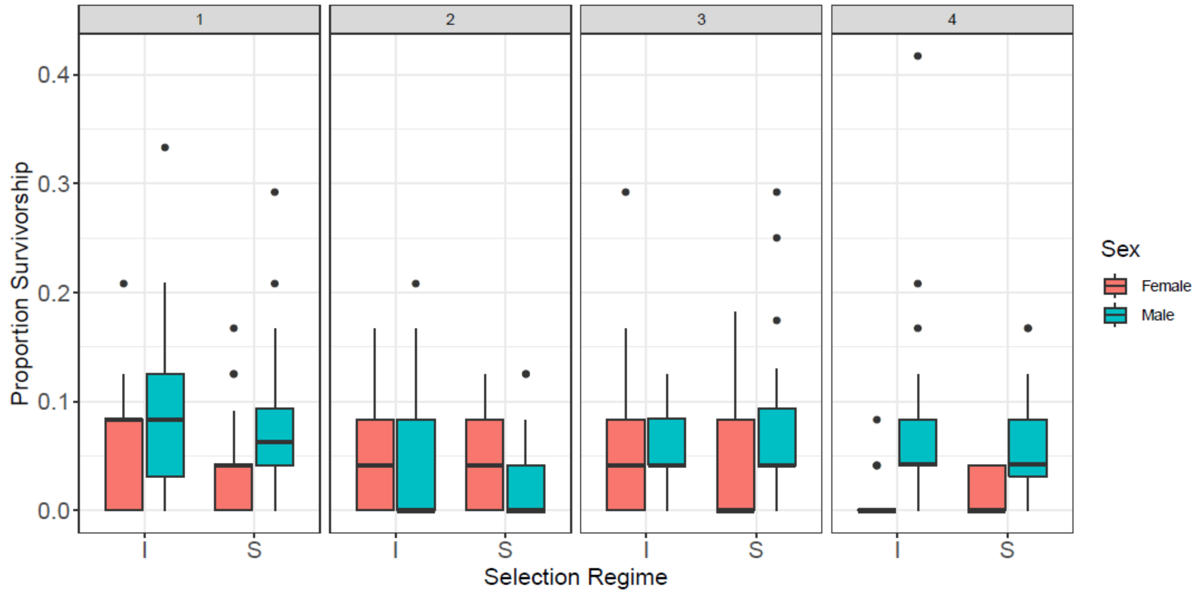


**Figure S2A** Effect of selection regime and sex on proportion survivorship at the end of the observation window for the X-Cloning Experiment

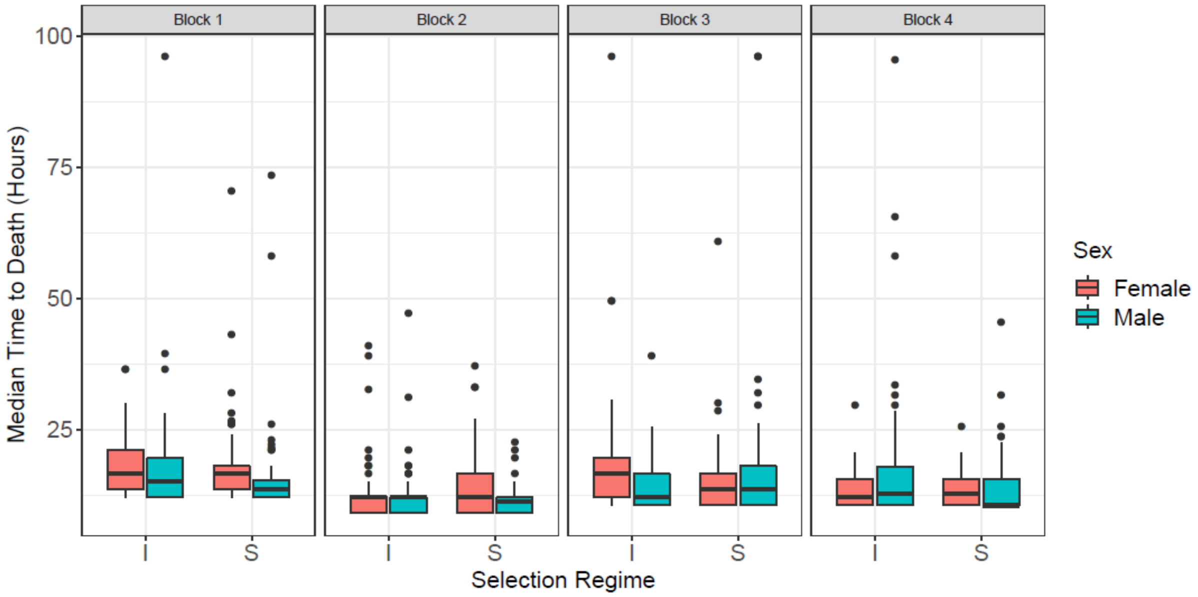


**Figure S2B** Effect of selection regime and sex on median time to death in the X-Cloning Experiment.

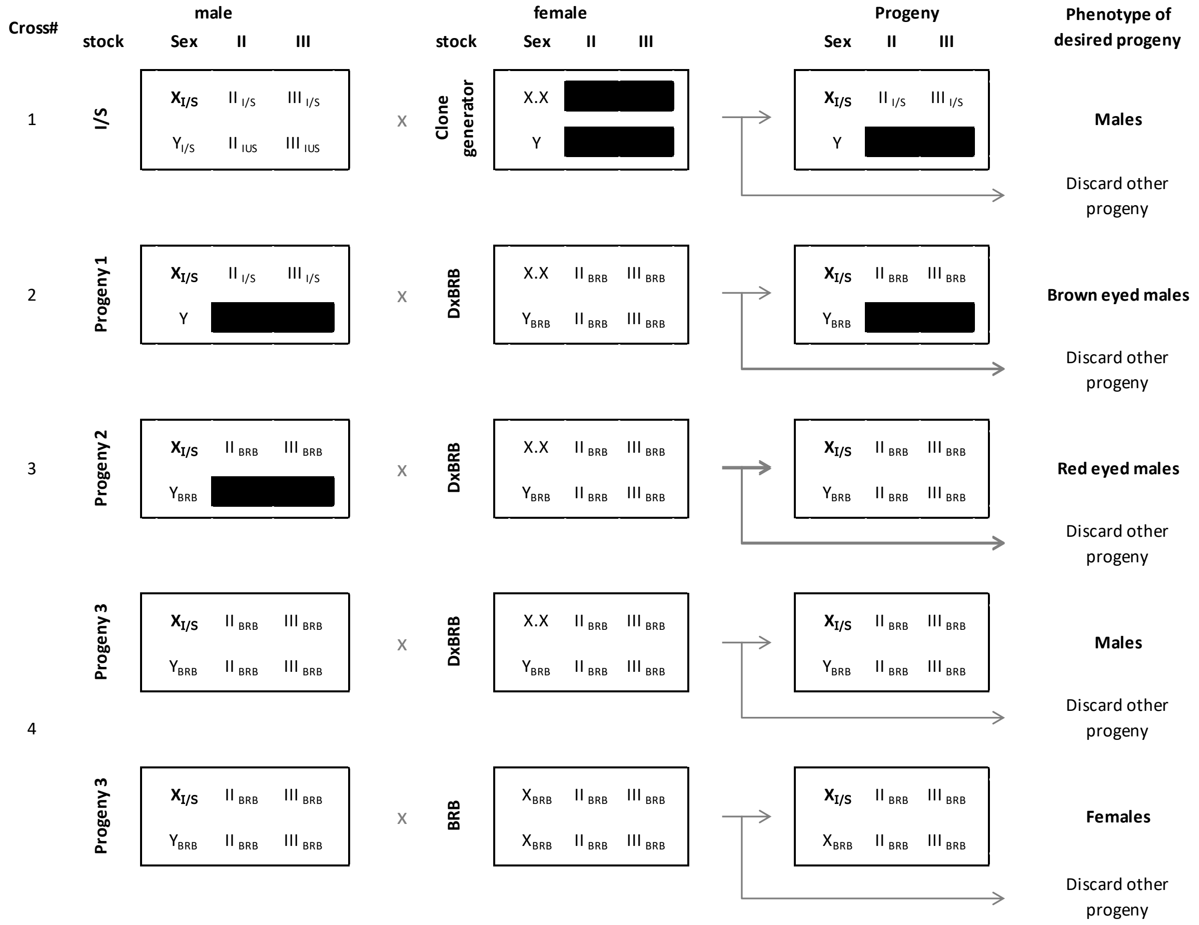


**Figure S3** A schematic for the crossing scheme used in the X-Cloning Experiment. The translocated 2nd and 3rd chromosomes are represented by a black bar since they must be inherited together. X, Y, II and III represent the respective chromosomes. X.X represents the compound X chromosome. The subscript represents the fly stock/selection regime from which the chromosome originates.

**Additional Analyses:**

We calculated the average median time to death and proportion survivorship for each X-line in both the sexes. Average median time to death was calculated as the mean of MTTD across the all vials of an X-line.

For these two read-outs of immunity, we calculated the correlation between male and female immunity.

We fit the following linear model separately for each combination of selection regime and block:

FemaleMeasure ~ MaleMeasure

We also calculated correlation using Spearman's Rank Correlation.

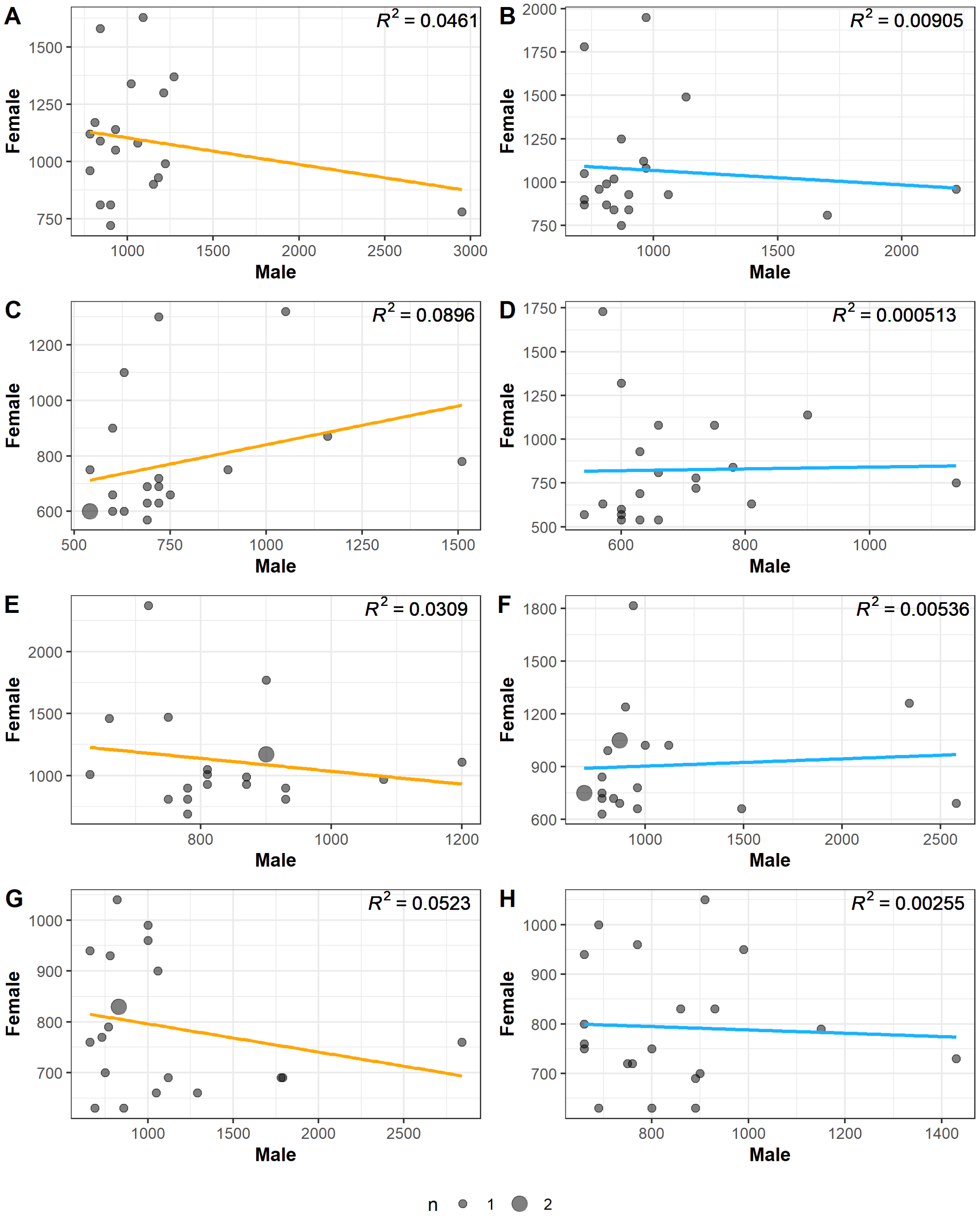


**Figure S4A.** Correlation between median time to death of males and females from the same X lines in A) I1, B) S1, C) I2, D) S2, E) I3, F) S3, G) I4, H) S4

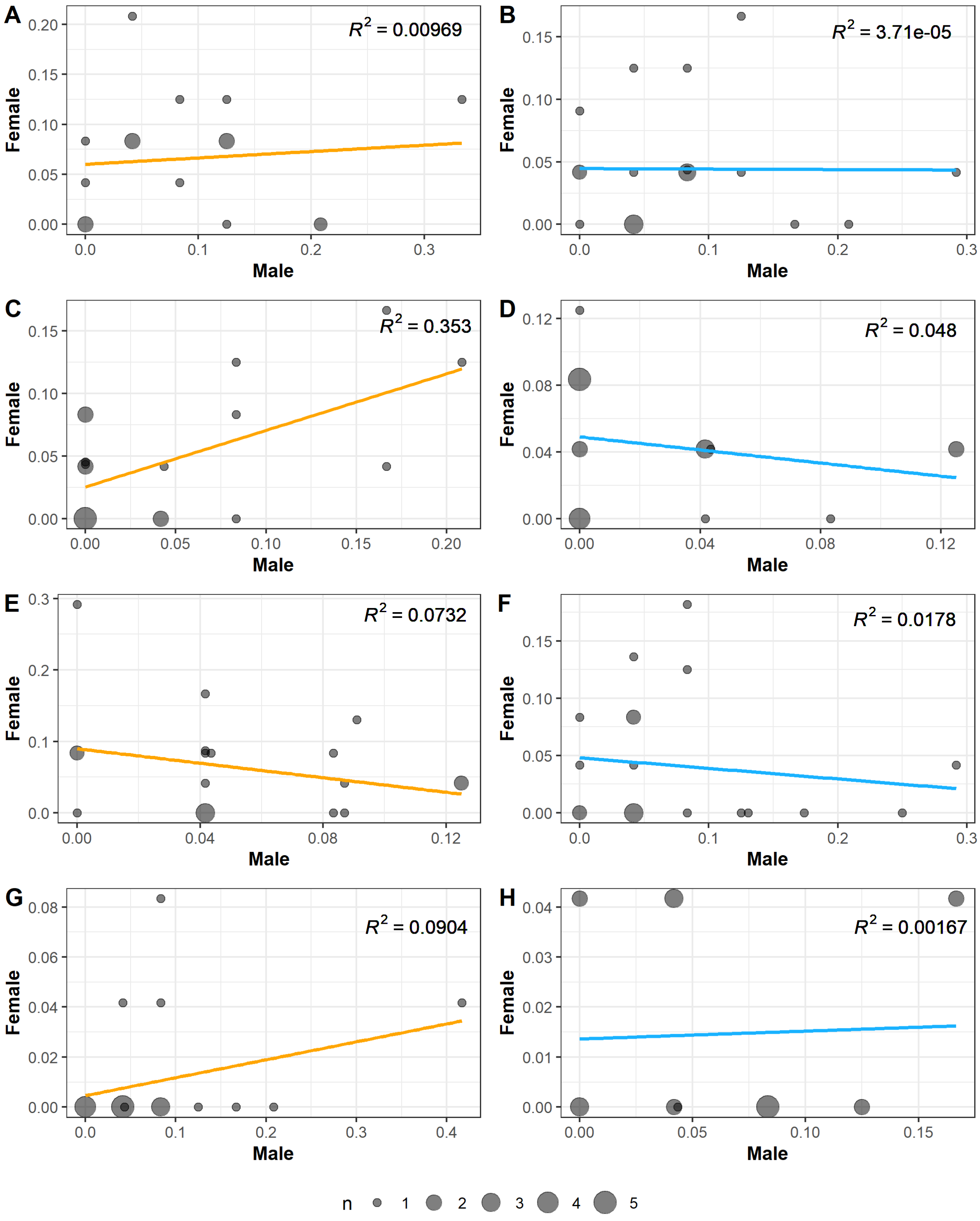


**Figure S4B.** Correlation between proportion survivorship of males and females from the same X lines in A) I1, B) S1, C) I2, D) S2, E) I3, F) S3, G) I4, H) S4

**Table S2A.** Results of correlation analysis between median time to death of males and females of the same X line by A) Linear Model and B) Spearman's rank correlation

**A) Linear Model**

| | | Estimate | Std. Error | t value | p value |
|---|---|---|---|---|---|
| I1 | (Intercept) | 1219.7436 | 151.8339 | 8.0330 | **<0.0001** |
| | Male | -0.1162 | 0.1281 | -0.9070 | 0.3770 |
| I2 | (Intercept) | 561.9780 | 164.5289 | 3.4160 | **0.0031** |
| | Male | 0.2787 | 0.2094 | 1.3310 | 0.1998 |
| I3 | (Intercept) | 1554.2376 | 584.9196 | 2.6570 | **0.0160** |
| | Male | -0.5193 | 0.6858 | -0.7570 | 0.4590 |
| I4 | (Intercept) | 851.7619 | 66.0325 | 12.8990 | **<0.0001** |
| | Male | -0.0556 | 0.0558 | -0.9970 | 0.3320 |
| S1 | (Intercept) | 1150.9281 | 213.7533 | 5.3840 | **<0.0001** |
| | Male | -0.0835 | 0.2059 | -0.4050 | 0.6900 |
| S2 | (Intercept) | 789.7930 | 368.2072 | 2.1450 | **0.0459** |
| | Male | 0.0504 | 0.5246 | 0.0960 | 0.9245 |
| S3 | (Intercept) | 860.8082 | 154.4088 | 5.5750 | **<0.0001** |
| | Male | 0.0413 | 0.1325 | 0.3120 | 0.7590 |
| S4 | (Intercept) | 821.3178 | 135.2469 | 6.0730 | **<0.0001** |
| | Male | -0.0336 | 0.1568 | -0.2140 | 0.8330 |

**B) Spearman's Rank Correlation**

| | S | Rho | p value |
|---|---|---|---|
| I1 | 1142.5 | -0.0022 | 0.9929 |
| I2 | 722.04 | 0.4571 | 0.0427 |
| I3 | 1430.9 | -0.0758 | 0.7506 |
| I4 | 1643.7 | -0.2358 | 0.3168 |
| S1 | 1255 | 0.0564 | 0.8132 |
| S2 | 987.65 | 0.2574 | 0.2732 |
| S3 | 1098.8 | 0.1738 | 0.4636 |
| S4 | 1321.9 | 0.0061 | 0.9798 |

**Table S2B.** Results of correlation analysis between proportion suvivorship of males and females of the same X line by A) Linear Model and B) Spearman's rank correlation

**A) Linear Model**

| | | Estimate | Std. Error | t value | p value |
|---|---|---|---|---|---|
| I1 | (Intercept) | 0.0601 | 0.0195 | 3.0730 | **0.0069** |
| | Male | 0.0638 | 0.1564 | 0.4080 | 0.6884 |

| | | | | | |
|---|---|---|---|---|---|
| I2 | (Intercept) | 0.0253 | 0.0114 | 2.2140 | **0.0400** |
| | Male | 0.4529 | 0.1446 | 3.1330 | **0.0058** |
| I3 | (Intercept) | 0.0898 | 0.0277 | 3.2470 | **0.0045** |
| | Male | -0.5081 | 0.4261 | -1.1920 | 0.2486 |
| I4 | (Intercept) | 0.0046 | 0.0067 | 0.6880 | **0.5000** |
| | Male | 0.0718 | 0.0537 | 1.3370 | 0.1980 |
| S1 | (Intercept) | 0.0445 | 0.0163 | 2.7360 | **0.0136** |
| | Male | -0.0039 | 0.1509 | -0.0260 | 0.9797 |
| S2 | (Intercept) | 0.0491 | 0.0100 | 4.9080 | **0.0001** |
| | Male | -0.1965 | 0.2063 | -0.9530 | 0.3533 |
| S3 | (Intercept) | 0.0481 | 0.0179 | 2.6810 | **0.0153** |
| | Male | -0.0925 | 0.1620 | -0.5710 | 0.5752 |
| S4 | (Intercept) | 0.0136 | 0.0073 | 1.8540 | 0.0802 |
| | Male | 0.0157 | 0.0903 | 0.1730 | 0.8642 |

**B) Spearman's Rank Correlation**

| | S | Rho | p value |
|---|---|---|---|
| I1 | 934.09 | 0.1806 | 0.4593 |
| I2 | 810.33 | 0.3907 | 0.0885 |
| I3 | 1486.8 | -0.1179 | 0.6206 |
| I4 | 932.02 | 0.2992 | 0.2000 |
| S1 | 1232.9 | 0.0730 | 0.7597 |
| S2 | 1657.3 | -0.2461 | 0.2955 |
| S3 | 1463.5 | -0.1004 | 0.6736 |
| S4 | 1441.4 | -0.0838 | 0.7254 |

We fit the following cox’s proportional hazards model separately for each block:

Time to Death ~ SelectionRegime + Sex + SelectionRegime:Sex + (1 | Infector) + (1 | SelectionRegime/Xline)

**Table S3.** Summary of blockwise analysis of survivorship post-infection by Cox Proportional Hazards model

**Cox Proportional Hazards**

**Block 1**

| Fixed Coefficients | coef | se(coef) | z value | p value |
|---|---|---|---|---|
| SelectionS | 0.0952 | 0.0953 | 1 | 0.3200 |
| Sexmale | 0.0246 | 0.0691 | 0.36 | 0.7200 |
| SelectionS:Sexmale | 0.0907 | 0.0970 | 0.93 | 0.3500 |
| Random effects | Variance | | | |
| Selection/Xline | 0.0456 | | | |

| | | | | |
|---|---|---|---|---|
| Selection | <0.0001 | | | |
| Infector | 0.0044 | | | |
| **Block 2** | | | | |
| Fixed Coefficients | coef | se(coef) | z value | p value |
| SelectionS | -0.0605 | 0.1280 | -0.47 | 0.6400 |
| Sexmale | 0.0849 | 0.0678 | 1.25 | 0.2100 |
| SelectionS:Sexmale | 0.1789 | 0.0961 | 1.86 | 0.0630 |
| Random effects | Variance | | | |
| Selection/Xline | 0.1157 | | | |
| Selection | 0.0002 | | | |
| Infector | 0.0062 | | | |
| **Block 3** | | | | |
| Fixed Coefficients | coef | se(coef) | z value | p value |
| SelectionS | 0.2444 | 0.0946 | 2.58 | **0.0097** |
| Sexmale | 0.2315 | 0.0677 | 3.42 | **0.0006** |
| SelectionS:Sexmale | -0.3304 | 0.0963 | -3.43 | **0.0006** |
| Random effects | Variance | | | |
| Selection/Xline | 0.0418 | | | |
| Selection | 0.0001 | | | |
| Infector | 0.1706 | | | |
| **Block 4** | | | | |
| Fixed Coefficients | coef | se(coef) | z value | p value |
| SelectionS | -0.0219 | 0.0953 | -0.23 | 0.8200 |
| Sexmale | -0.2665 | 0.0679 | -3.92 | **<0.0001** |
| SelectionS:Sexmale | 0.1465 | 0.0951 | 1.54 | 0.1200 |
| Random effects | Variance | | | |
| Selection/Xline | 0.0461 | | | |
| Selection | <0.0001 | | | |
| Infector | 0.1461 | | | |
